# MolAR: memory-safe library for analysis of MD simulations written in Rust

**DOI:** 10.1101/2024.09.21.614237

**Authors:** Semen Yesylevskyy

## Abstract

Transition to the memory safe natively compiled programming languages is a major software development trend in recent years, which eliminates memory-related security exploits, enables a fearless concurrency and parallelization, and drastically improves ergonomics and speed of software development. Modern memory-safe programing languages, such as Rust, are currently not used for developing molecular modeling and simulation software despite such obvious benefits as faster development cycle, better performance and smaller amount of bugs. In this work we present MolAR -the first memory-safe library for analysis of MD simulations written in Rust. MolAR is intended to explore the advantages and challenges of implementing molecular analysis software in the memory-safe natively compiled language and to develop specific memory-safe abstractions for this kind of software. MolAR demonstrates an excellent performance in benchmarks outperforming popular molecular analysis libraries, which makes it attractive for implementing computationally intensive analysis tasks.

MolAR is freely available under Artistic License 2.0 at https://github.com/yesint/molar.

## Introduction

Memory safety has become a major software development trend in recent years due to the pressing issues imposed by traditional programming languages and development practices ^1^. Traditionally the niche of high-performance compiled programming languages was dominated by C and C++, which were deliberately designed to give the programmer full control on memory management and did not contain built-in guarantees of safe memory access. While arguably useful for the system-level programming, such as creation of operating system kernels and device drivers, this inherent memory unsafety has become an issue for applied programming in several different aspects. First, it leads to a plethora of memory-related bugs, such as buffer overflows, use of invalid pointers, memory leaks and corruption, which are often utilized by security exploits ^2^. Second, it greatly hampers adoption of concurrency and multithreading due to the uncontrollable race conditions, which make parallel programs notoriously difficult to write and debug. Third, it severely affects the ergonomics of software development and imposes a high entry barrier for beginners and non-professional programmers, such as researchers and PhD students.

Currently we witness a new generation of programming languages that are built around memory safety as the core concept. The most mature and widespread of them is Rust, which is currently considered one of the most promising alternatives to C and C++. Rust ensures memory safety on the level of type system by means of strict built-in ownership and borrowing rules ^3,4^. It already demonstrated impressive effectiveness in multiple domains from device drivers to high-load web servers and game engines.

The software for analysis of molecular dynamics simulations can also benefit from adopting the memory safe languages like rust. Modern MD simulations produce very large trajectories, which require significant time for processing and analysis. The analysis algorithms are often quite expensive computationally and it is not uncommon that the analysis takes comparable time of even longer than the production MD simulations themselves. Most of the analysis tasks could also benefit a lot from parallelization because they operate on individual trajectory frames independently. All this encourages implementation of the analysis software in the natively compiled languages. However, implementing them in C or C++ is tedious, slow and error-prone because of well-known shortcomings of these languages themselves and their corresponding ecosystems. The poor ergonomics of programming in memory unsafe languages in general and notorious complexity of concurrent programming in them in particular, are among the major factors, which discourages the community from using natively compiled languages for trajectory analysis.

The vast majority of modern software for the analysis of molecular dynamics simulations is written in Python. This is currently a de-facto requirement for a wide adoption in the simulation community because most of the researchers and students are familiar with this language. However, Python is one of the slowest among popular scripting languages ^5,6^. It is also poorly suitable for parallelization and multithreading due to the Global Interpreter Lock mechanism built into its interpreter ^7^. As a result such popular libraries as MDAnalysis ^8,9^ and MDTraj ^10^, which are implemented mostly in Python, are rather slow despite calling C and C++ libraries under the hood for performance-critical operations like reading trajectory files and performing matrix arithmetics. This is clearly visible when compared to Pteros ^11,12^, which is written in C++ and provides only a thin Python wrapper that directly calls optimized compiled backend.

Pteros is currently the only actively developed open source molecular analysis library, which is implemented in a natively compiled language (C++). However, for more than fifteen years of existence it did not attract enough contributors, which is, to a large extent, a consequence of the poor reputation of C++ in the simulation community as an “overcomplicated legacy language”. Memory unsafety of C++ has also manifested itself multiple times during the Pteros development cycle and introduced a number of hard to detect bugs, especially in the parallel code.

Thus there is a clear demand for a new natively compiled molecular analysis library, which would be faster than existing Python solutions, will allow straightforward parallelization and will be in line with modern memory safety trends.

Here we present MolAR -the Molecular Analysis Library for Rust. MolAR is a research project written in Rust, which explores the advantages and challenges of implementing the general-purpose toolbox for analysis of MD trajectories in the memory-safe natively compiled language. In this paper we cover design and implementation of the cornerstone data types for molecular analysis within the Rust type system and ownership rules. Then we discuss the technical details of implementation and provide the comparison of performance between MolAR, Pteros, MDAnalysis, MDTraj and Gromacs analysis tools.

## Memory-safe abstractions for molecular analysis

### Terminology

All libraries for analysis of MD simulations are built around three major entities: *Topology, State* and *Selection*. These entities may have different names or could be encapsulated into the higher-level abstractions, but they are required to properly create, read, query and modify the systems of atoms originating in MD simulations.

- *Topology* contains the time-independent system properties, such as identity of the simulated atoms (type, name, charge, mass, etc.), information about the residues, chains and molecules, bonds, etc.
- *State* contains information, which changes in the course of simulation: positions, velocities and forces of the atoms, simulation periodic box, simulation time and step number, etc.
- *Selection* is a view into Topology and concrete State, which points to a subset of atoms. Internally selections are usually implemented as arrays of indexes or as the boolean mask arrays. Selections do not hold any data themselves but rather act as handles for manipulating the groups of atoms.

In Rust terms selections are mutable references into the underlying data stored in *Topology* and *State* and as such they are subject to strict ownership and borrowing rules. There is a clear distinction in Rust between the data ownership (entity which is responsible for allocating and freeing the memory) and data borrowing (entities that temporarily reference the data, but are not managing its memory). There are two types of borrowing: immutable (shared read-only access) and mutable (exclusive read-write access). For any object at most one mutable borrow or any number of immutable borrows may exist at any point in time. This is strictly enforced by the Rust compiler and is a cornerstone of Rust’s strong memory safety guarantees.

### Restrictions of Rust borrowing rules

Unfortunately, Rust borrowing rules do not play nicely with the concept of atom selections. Selections are inherently mutable (allow to modify properties of the referenced atoms) and may overlap arbitrarily (for example, the same Cα atom could be shared by selections for the protein backbone, particular residue and particular chain). The changes made by one selection are immediately visible by the others since they point to the same physical memory location. In Rust this type of memory access is often called an “aliased mutable borrowing” and is forbidden by design because it is not memory safe in the general case.

Although it is possible to design an API for atom selections in safe Rust by deferring “binding” of selection to the concrete *Topology* and *State* to the point of actual selection usage in the client code, the ergonomics of such a solution appears to be very poor. The user should pass around three objects (unbound selection, topology and state) instead of one (bound selection), the thread safety becomes notoriously hard to achieve and the life times of all three objects become logically separated, which makes reasoning about data flow even more complicated.

The analysis of existing strategies of addressing this issue shows that there are no standard mechanisms of achieving required level of ergonomics and safe memory access by selections that may reference the data from multiple threads. Although some concrete cases of the required mutable mutable aliasing could be made memory safe, the Rust borrow checker can not detect such patterns. However, the programmer can work around this limitation by using unsafe subset of the language and by exposing a safe API to the library users. This approach is considered idiomatic and is used heavily in Rust standard library and the popular packages. MolAR uses this approach to provide safe selection objects, which use unsafe operation under the hood to manage the multiple mutable aliasing.

### Memory-safe selection types

We analyzed typical selection usage patterns and classified them into four classes summarized in Table 1.

| Selection kind | Data access | May overlap? | Require bounds checking |
| --- | --- | --- | --- |
| MutableSerial | read / write | yes | no |
| BuilderSerial | read / write / | yes | yes |
| ImmutableParallel | read only | yes | no |
| MutableParallel | read / write | no | no |

In order to account for these selection kinds we decided to use a single generic type *Sel<K>* parameterized by the zero-size marker type K, which holds all necessary information about selection behavior.

We also want to guarantee statically at compile time that a given pair of *Topology* and *State* could only be used to create selection of just one particular kind at a time. This simplifies reasoning about data access a lot and does not impose any significant usability constraints. We enforce such a behavior by wrapping *Topology* and *State* into the auxiliarily data structure called *Source*, which is parameterized by the marker types K and produces selections of kind K. *Topology* and *State* are moved into the *Source* during its construction and selection could only be created by the methods of the *Source* itself.

Let us briefly discuss each of the selection kinds. *MutableSerial* selections are mostly equivalent to selections familiar from other molecular modeling libraries. They are only used from the same thread that created them and are not allowed to be shared between the threads. Since all operations on such selections are sequential by definition, no race conditions may occur and it is safe to have multiple aliasing mutable references to the underlying data (Fig 1).

**Figure 1.**
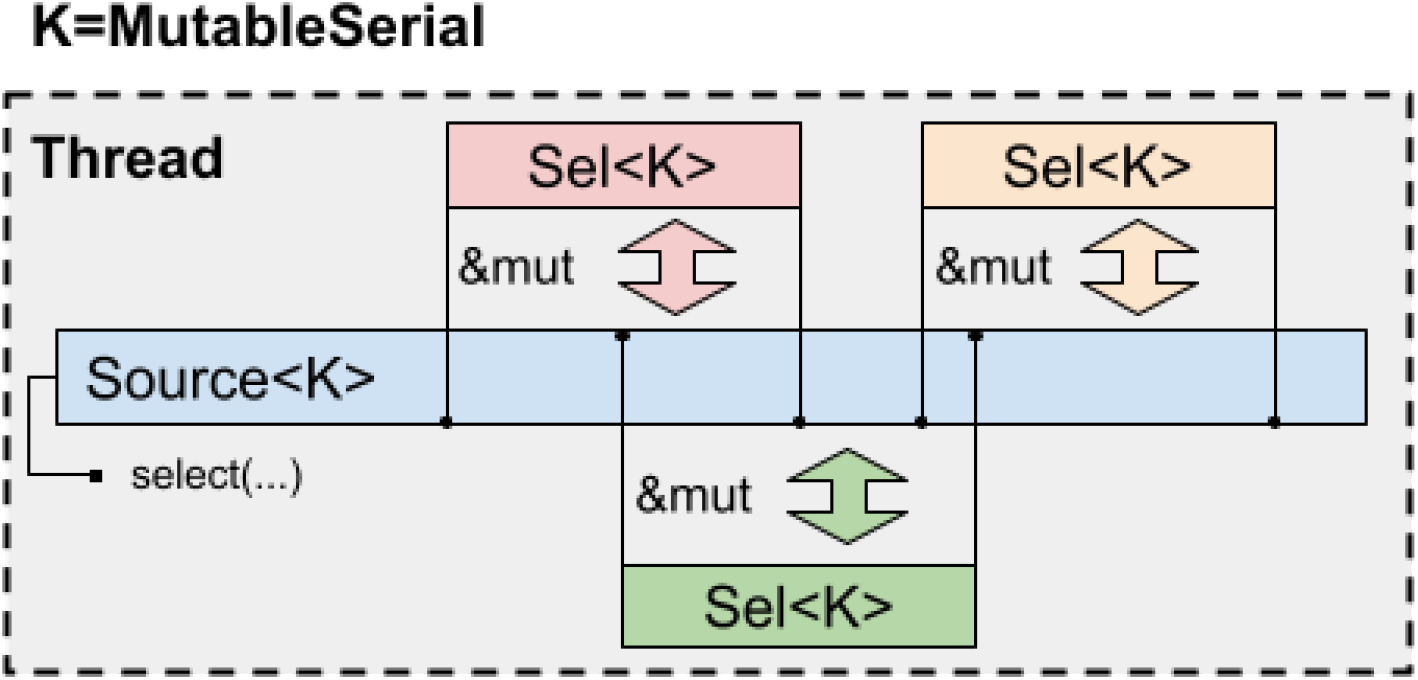
Scheme of data access for MutableSerial selections. On this and subsequent figures, for the sake of clarity, selections are shown as referring to contiguous spans of indexes in the underlying data. In reality selections may refer to arbitrary non-contiguous sets of indexes.

There is also a special case of serial selections (*BuilderSerial*) that reference *Topology* and *State* with variable number of atoms. This is common during the initial building of simulation systems or when addition or deletion of auxiliary dummy or virtual particles is required for analysis. In this case selections should additionally check whether they refer to the atoms that exist in the system to avoid invalid memory access after deleting some atoms. For other kinds of selections such bound checking is not needed since it is guaranteed that the number of atoms does not change during their lifetime. To ensure this only the *Source<BuilderSerial>* objects implement the methods to add or delete atoms (Fig. 2).

**Figure 2.**
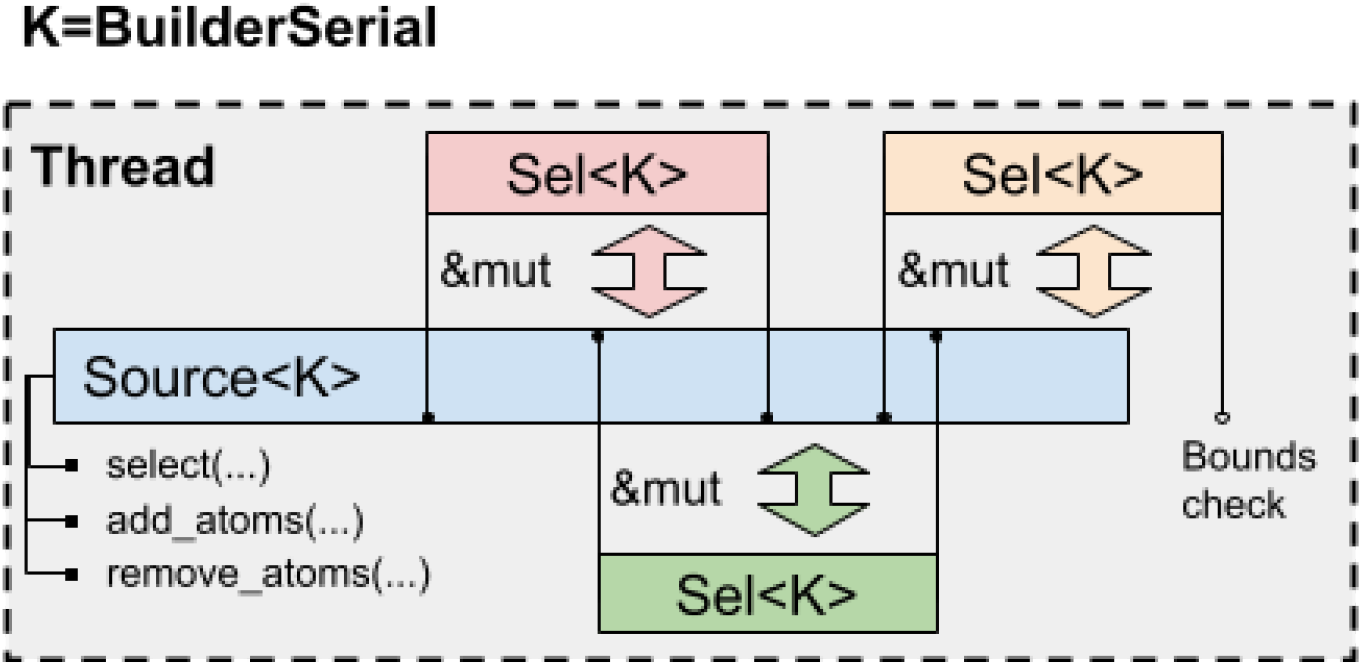
Scheme of data access for BuilderSerial selections.

Bound checking is implemented through the methods of the marker type *K*=*BuilderSerial*. For all other types *K* that do not require bound checking an empty method is implemented, which is eliminated by the optimizing compiler. In such a way there is no runtime cost for the selection that does not require this feature.

In other libraries there is no distinction between *MutableSerial* and *BuilderSerial* selection and thus the bound checks are either implemented for all of them or not implemented at all ignoring possible invalid memory access.

Parallel selections are intended to be used from multiple threads while referencing shared *Topology* and *State* in a safe way without data races. There are no such abstractions in the majority of other molecular analysis libraries. Behavior of parallel selections depends on whether or not they may overlap.

The *MutableParallel* selection kind guarantees that selections do not overlap and thus it is safe to hold multiple aliasing mutable references to the underlying data despite Rust borrow checker is unable to deduce that they refer to different distinct array elements (Fig. 3). The non-overlapping of selections is guaranteed by the *Source*<*MutableParallel*>, which is used to create them. Internally it contains a hash set of used atom indexes and each new selection is checked against it. This run time cost should be paid only once during creation of selections while using them is free from any run time overhead.

**Figure 3.**
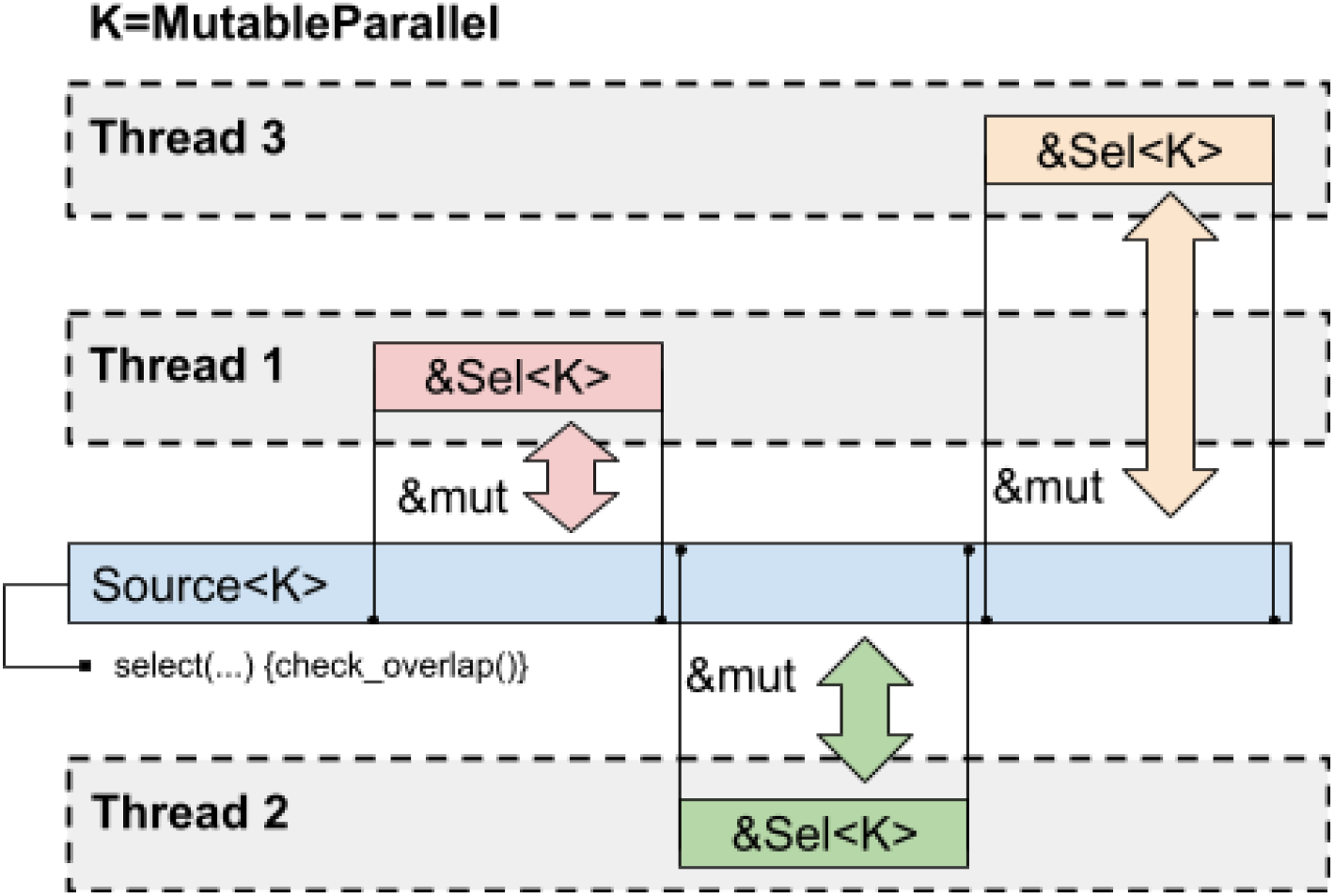
Scheme of data access for MutableParallel selections.

If parallel selections may overlap then it is only safe to have multiple immutable read-only references to the underlying data. This access pattern is implemented by the *ImmutableParallel* selections that only implement those traits that provide read-only access to the data (Fig. 4). No overlap check is required in this case.

**Figure 4.**
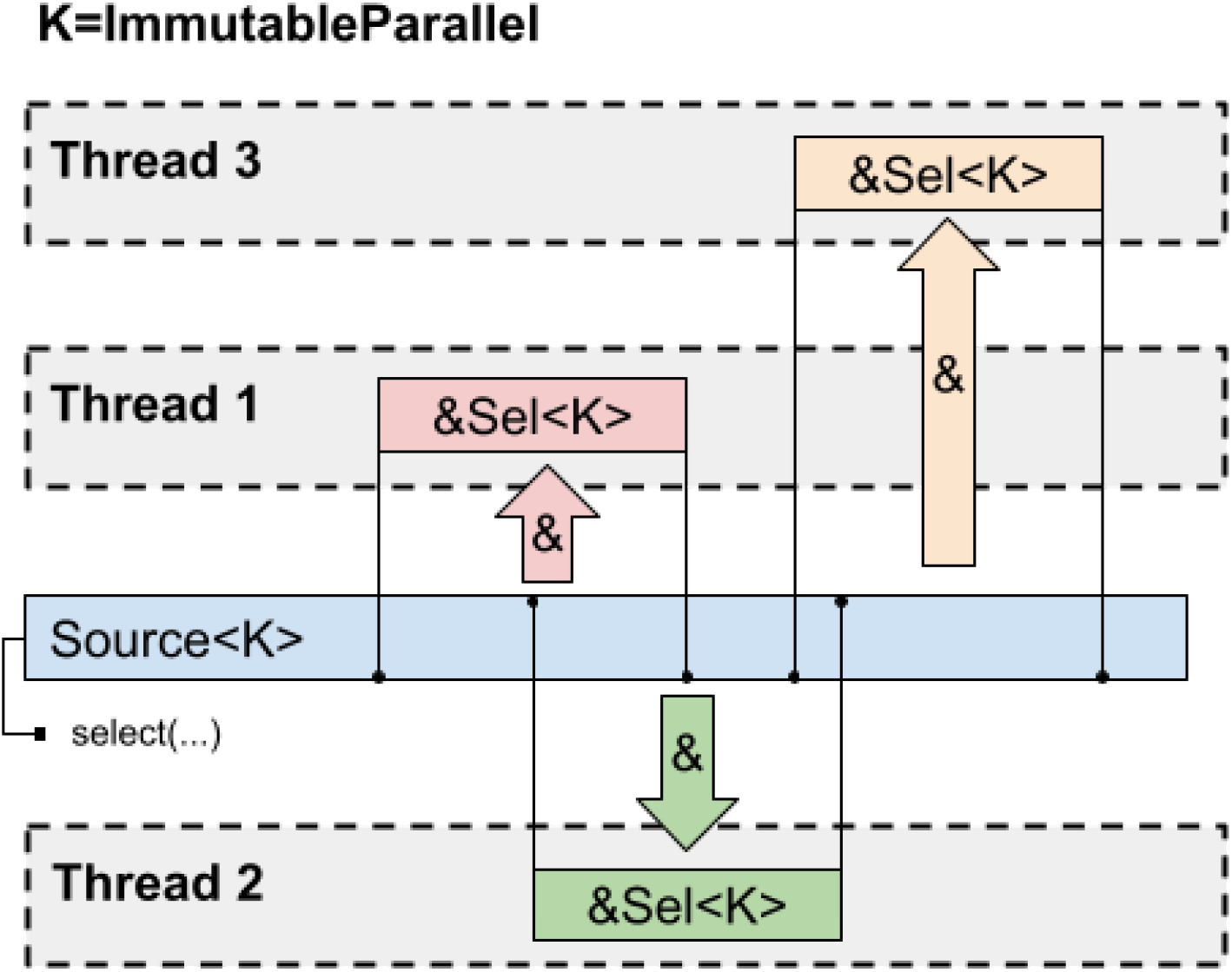
Scheme of data access for ImmutableParallel selections.

### Sub-selections

One of the useful features that was introduced in Pteros and which is often missing in the other analysis libraries, is an ability to create sub-selections by narrowing down already existing selections. Sub-selections are much faster to create since they only require looping through the indexes of existing selection instead of the whole system. They also facilitate flexible convenience methods for splitting selections in different ways (for example, into individual residues or chains).

Sub-selections are trivial to implement for all selection kinds except the *MutableParallel*. Indeed, *MutableParallel* selections should never overlap, while any sub-selection that exists at the same time as the parent selection will violate this requirement. In order to deal with this issue MolAR introduces three distinct ways of creating sub-selections: subselecting, splitting and fragmentation (Table 2).

**Table 2.**
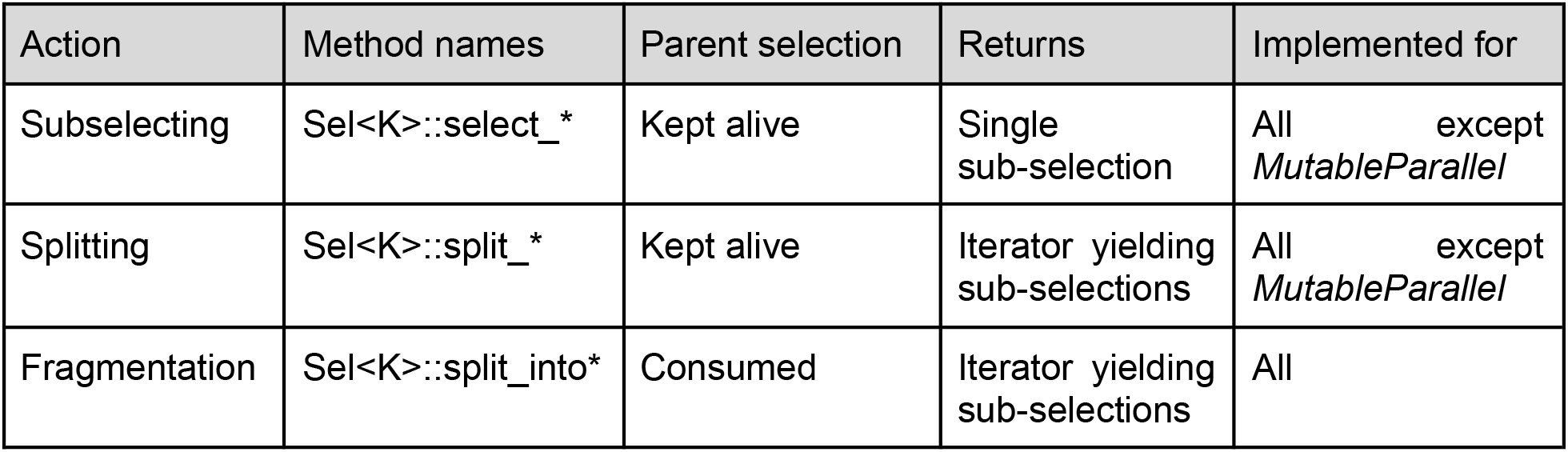
Characteristics of different techniques of creating sub-selections.

| Action | Method names | Parent selection | Returns | Implemented for |
| --- | --- | --- | --- | --- |
| Subselecting | Sel<K>::select_* | Kept alive | Single sub-selection | All except *MutableParallel* |
| Splitting | Sel<K>::split_* | Kept alive | Iterator yielding sub-selections | All except *MutableParallel* |
| Fragmentation | Sel<K>::split_into* | Consumed | Iterator yielding sub-selections | All |

Subselecting creates a single new selection while keeping the parent one alive. Splitting splits the parent selection into multiple non-overlapping sub-selections using user-provided splitting logic while keeping the parent one alive. Fragmentation is the same as splitting but it consumes the parent selection, which guarantees that there is no overlap between the sub-selections and their parent. *MutableParallel* selections only implement fragmentation, while all other selection kinds implement all three techniques. Figure 5 illustrates the memory layout of sub-selections originating from different techniques.

**Figure 5.**
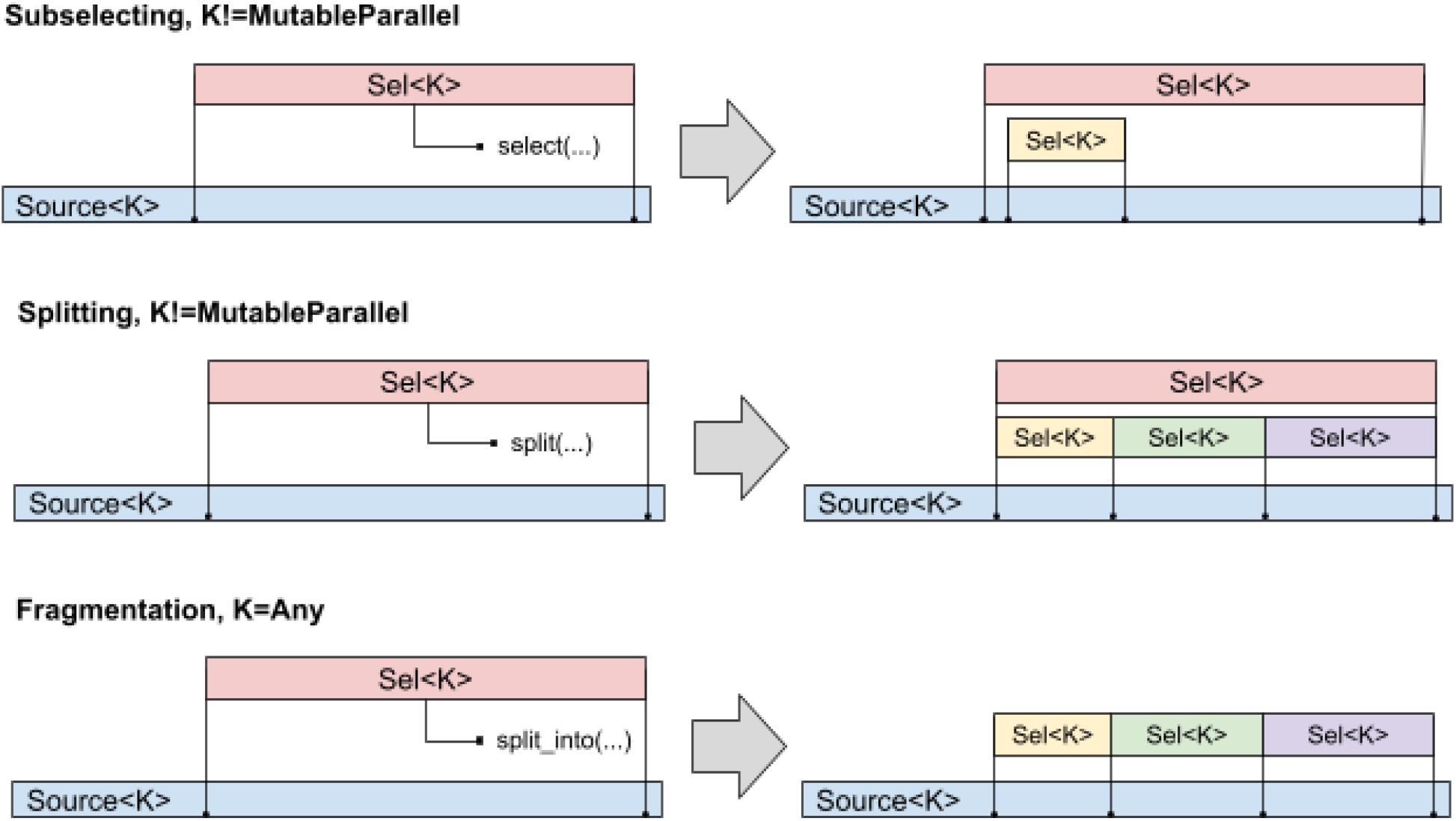
Schematic memory layouts created by different techniques of creating sub-selections.

### Analysis and IO traits

One of the strong points of Rust is the concept of traits, which define the behavior that could be shared by multiple types and are checked at compile time. Selective implementation of the traits allows for fine-grained control of types’ behavior and also allows to separate data manipulations from the higher-level logic.

MolAR defines three categories of traits, which are used for IO operations, read-only and read-write access to the atoms. The hierarchy of traits is rooted in general-purpose provider traits that expose concrete properties, mostly as iterators. For example, the *AtomsProvider* trait defines the *iter_atoms()* methods, which returns an iterator over the atoms contained in the object that implements it. *Topology* and *Source* will return an iterator over all atoms in the system, while *Sel* will provide an iterator over the selected atoms. Not all the properties are iterable, for example, the *BoxProvider* trait exposes the optional periodic box of the system.

The random-access traits allow addressing the properties by an atom index, which is required for algorithms that can not be implemented efficiently with the single-pass forward iterators only, such as removing the jumps over periodic box for covalently bound molecules.

### Technical details

#### Data layout

The actual data is stored in internal *TopologyStorage* and *StateStorage* types. They are wrapped into *SyncUnsafeCell* “magic” wrapper type, which instructs the Rust compiler that the data is not guaranteed to be uniquely mutably borrowed and thus is allowed to violate the borrow checker rules without confusing the optimizer and the low-level assembly code generation backend. This wrapper is also allowed to be shared between the threads, which is necessary for implementing parallel selections. *Topology* and *State* types are just wrapper over *TopologyStorage* and *StateStorage* that provide necessary semantics.

Upon construction, which typically happens by reading from file, *Topology* and *State* are wrapped into the atomic reference counted smart pointers *Arc* from the *triomphe* crate ^13^. Rust standard *Arc* smart pointers are less convenient for our goals and also contain the weak pointer counter, which is not needed in our use case.

*Triomphe* crate provides a *UniqueArc* pointer that is guaranteed to be uniquely owned and could be then converted to conventional *Arc* shared pointer. When the *Source* is created the *UniqueArc<Topology>* and *UniqueArc<State>* pointers are expected while the corresponding *Arc* pointers are stored internally. This prevents *Topology* and *State* from accidentally leaking and being used in multiple *Sources*.

When selections are created from a *Source* they also refer to *Topology* and *State* by means of *Arc* pointers. This guarantees that the lifetime of underlying data is always tied to the last selection, which stays alive. The lifecycle of smart pointers and the relations between different types is shown in Fig. 7.

**Figure 6.**
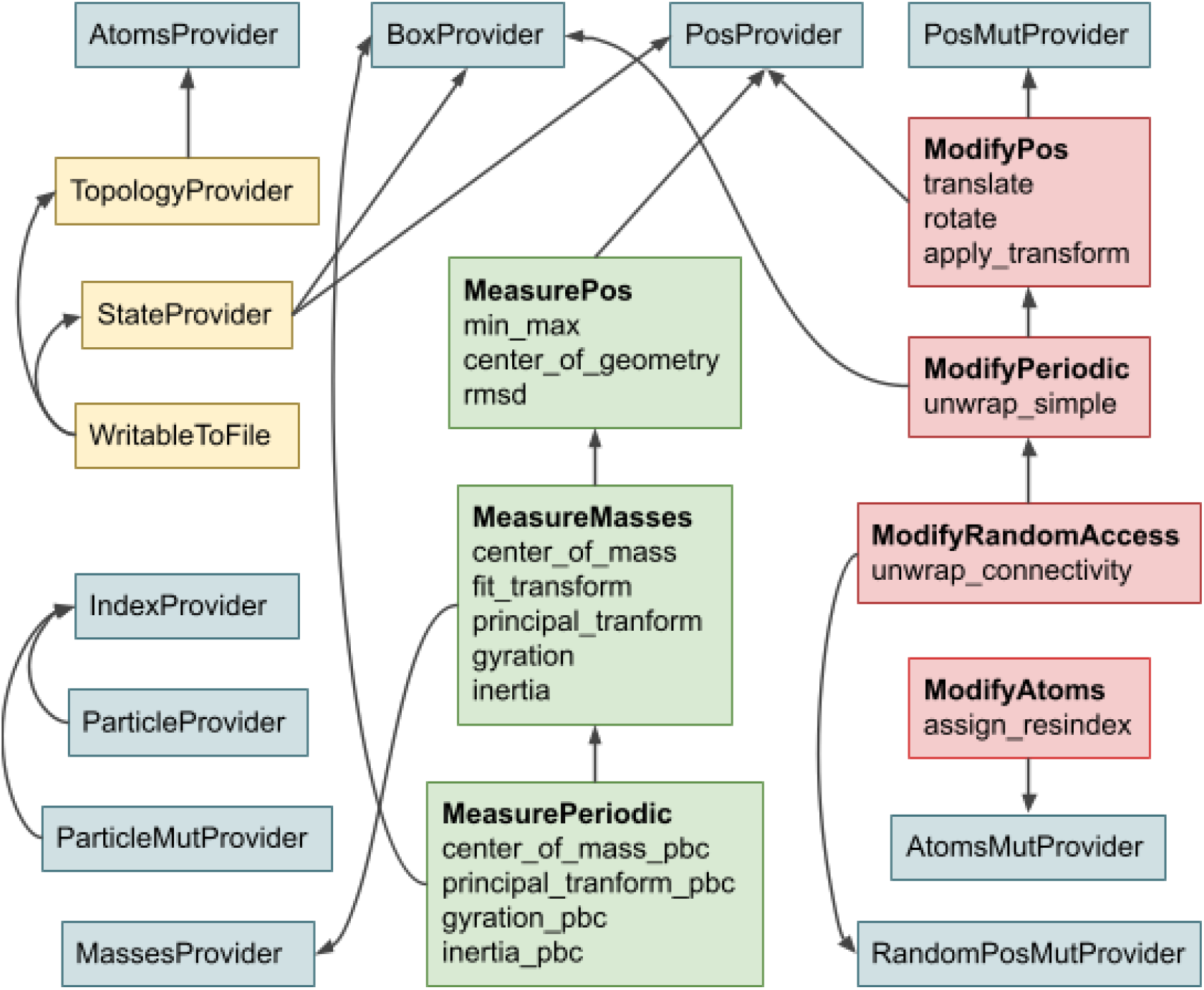
The hierarchy of main MolAR traits. The general purpose traits are blue, the IO traits are yellow, the read-only analysis traits are green (“measure traits”) and the read-write analysis traits (“modify traits”) are red. The arrows show trait dependencies. For analysis traits some of the provided methods are shown.

**Figure 7.**
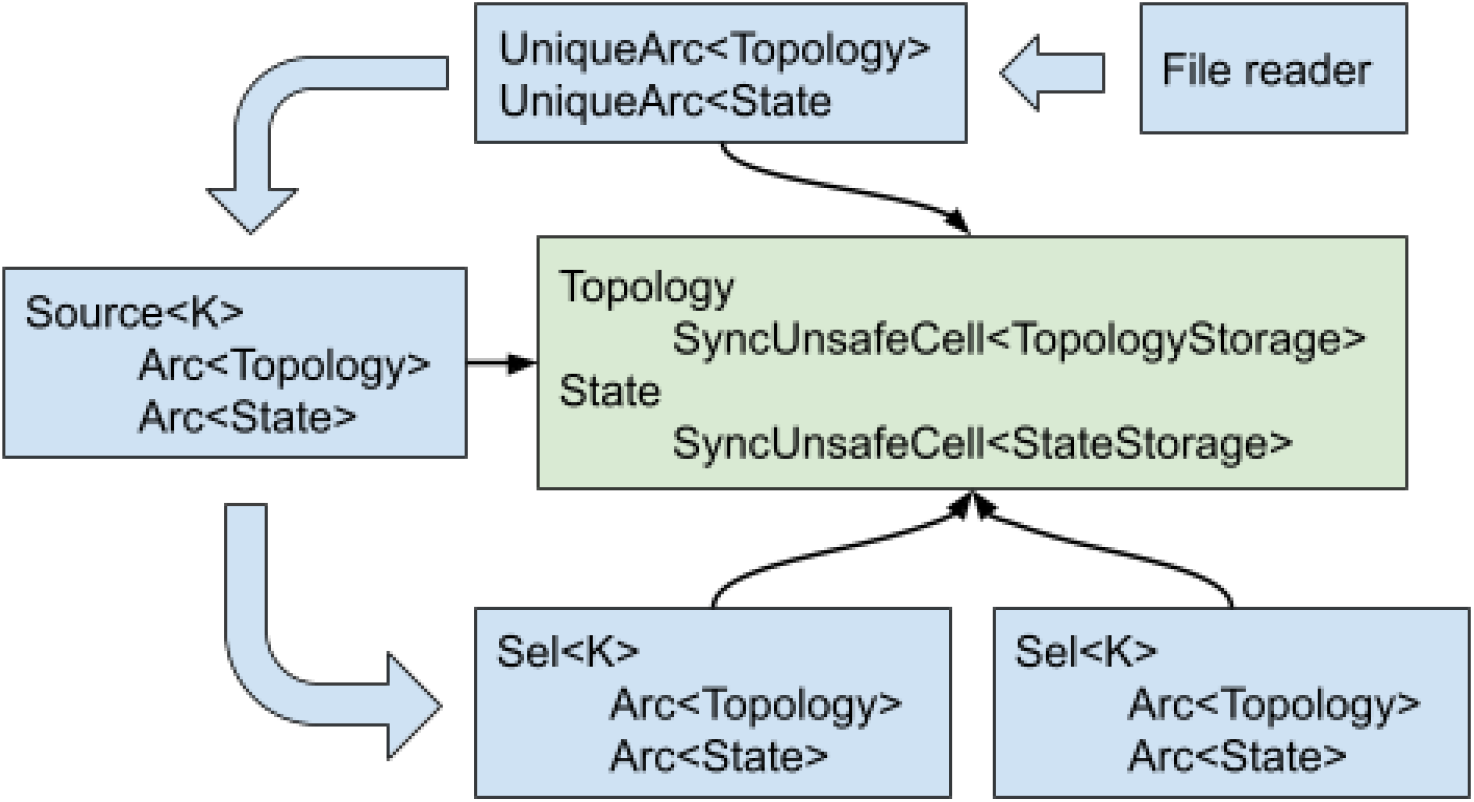
The scheme of relations between the MolAR data types and the life cycle of smart pointers holding Topology and State data. Wide arrows show transformations of the user-exposed data types (blue rectangles), while thin arrows show the internal pointers to actual raw data hidden from the user (green rectangles).

The marker types *K* used to parameterize selections and sources are also used to control their passing to different threads. Serial selections do not implement *Send* and *Sync* traits and thus are statically guaranteed to be only used in the same thread where they are created. The same is true for *Source*, which should always stay in the same thread that created it.

#### Support for file formats and third-party libraries

MolAR utilizes a pragmatic approach to reusing existing code written in C and C++ for performing low-level IO operations. The bindings to existing libraries are generated by the bindgen crate ^14^. The source code of third-party libraries is bundled with MolAR and is compiled together with the rust code by means of cmake crate ^15^. The only additional dependencies are the C/C++ compiler and the cmake itself.

MolAR relies on well-established low-level libraries for reading and writing the most popular molecular data formats. The bindings to VMD molfile plugins written in C are available in the auxiliary molar_molfile crate ^16^ and currently provide access to PDB, DCD and XYZ formats. Other formats supported by VMD molfile plugins could be added trivially in an incremental way.

Support for Gromacs XTC files is provided by the bindings to a customly modified xdrfile library encapsulated into molar_xdrfile crate ^17^. Modifications enable ergonomic random-access to trajectory frames, which greatly speeds up such operations as skipping initial unequilibrated part of the trajectory or extracting the final system state.

There is also a seamless support for reading Gromacs run input files in TPR format. This is implemented in the same way as in Pteros by linking to locally installed gromacs and using the necessary C header files from the matching Gromacs source tree. Unfortunately, Gromacs does not expose the functions for reading and parsing TPR files in its public API, so hacking into the private Gromacs headers is necessary. On the positive side such an approach accommodates automatically to any new Gromacs version since no custom parsing of the TPR files is performed and all processing is done by the Gromacs core library itself. This functionality is provided by the molar_gromacs crate ^18^.

There is also an optional feature of downloading and compiling the bare minimal version of Gromacs during MolAR compilation itself. In this case no external Gromacs installation is needed and the latest git version of Gromacs is used to ensure backward compatibility with all existing stable versions. Gromacs is a very heavy dependency, which drastically increases the compilation time, so this feature is switched off by default and is reserved for installing MolAR in a fully self-consistent way in such environments as docker containers.

MolAR also bundles a modified version of the powersasa C++ code ^19^ for computing the solvent accessible surface area (SASA).

## Benchmarks

The main motivation of creating MolAR is to provide a solid set of memory-safe abstractions for the analysis of MD trajectories. There are no specific performance-oriented optimizations in the MolAR code except the usage of unsafe access to array elements without the bound check in the cases where the indexes are guaranteed to be within the bounds by definition. Thus, it is interesting to test the performance of MolAR in real-world scenarios and to compare it with other trajectory analysis programs and libraries.

We compared MolAR with Pteros ^12^, Gromacs ^20^, MDAnalysis ^8,9^, MDTraj ^10^ and VMD ^21^. For benchmarks we have chosen a trajectory of a small protein with ∼4300 atoms and 4000 frames. We set up three benchmarks, which represent commonly occurring real-world analysis tasks (Table 3). First task is aligning protein with the reference structure in all frames and computing the RMSD for all trajectory frames.

**Table 3.** Comparison of different molecular analysis libraries and Gromacs suite of analysis programs in three real-world benchmark tasks. For MDAnalysis, MDTraj, Pteros and Molar the real execution time without program start up time is reported. For Gromacs the wall time including the startup time is reported. Time is reported in seconds.

| Benchmark: | On each frame align protein with reference and compute RMSD | On each frame compute center of masses of atoms within 1 nm from any atom of a given residue | Extract given selection into separate trajectory |
| --- | --- | --- | --- |
| MDAnalysis 2.7 | 5.10 ± 0.14 | 12.1 ± 0.19 | 1.0 ± 0.03 |
| MDTraj | 3.70 ± 0.19 | N/A | 0.79 ± 0.02 |
| VMD | 1.10 ± 0.3 | 0.70 ± 0.45 | 0.80 ± 0.4 |
| Gromacs 2023 | 0.68 ± 0.03 <sup>1</sup> | N/A | 0.58 ± 0.14 <sup>2</sup> |
| Pteros 2.5 gcc | 0.80 ± 0.005 | 2.64 ± 0.04 | 0.61 ± 0.003 |
| Pteros 2.5 clang | 0.66 ± 0.005 | 2.55 ± 0.02 | 0.57 ± 0.02 |
| Molar 0.4 | 0.61 ± 0.01 | 0.46 ± 0.007 | 0.47 ± 0.03 |

<sup>1</sup> The following command is used: `time echo 0 0 | gmx rms -f .aligned.xtc -s protein.pdb`
<sup>2</sup> The following command is used: `time echo 3 | gmx trjconv -f protein.xtc -s protein.pdb -o extracted.xtc`

Second task is selecting the atoms within 1 nm from any atoms of particular protein residue and computing the center of mass of this selection for all trajectory frames. There is no simple way to perform such an analysis in MDTraj and Gromacs, so we skipped them for this task.

The third task is reading the trajectory and writing another trajectory containing only a specified selection (a single residue). This is the most basic operation, which only requires trajectory reading and writing, so it is expected that the performance of all libraries will be comparable.

All tests were performed 10 times subsequently and the average run time is reported. The consumer laptop with AMD Ryzen 7 PRO 6850U CPU (16 logical cores) and 32 Gb of RAM running Fedora Linux 40 was used for testing. Pteros was compiled with GCC 14.2.1 and clang 18.1.8. MolAR was compiled with Rust 1.80.0. MDAnalysis and MDTraj were installed from the Python Package Index (PyPI) repository. Gromacs was compiled with GCC 14.2.1 and the VMD binary package for Linux was installed from the official site. All benchmarks were timed after initial cache warming to exclude the effect of caching disk IO operations.

Table 3 clearly shows that MolAR outperforms all other libraries by a significant margin for the first and second tasks. In contrast, the MDAnalysis, which is arguably the most popular trajectory analysis library at the time of writing, is the slowest of all tested techniques. Even more surprisingly, MDAnalysis is 1.8 times slower than MolAR even in the third task, which mostly involves IO operations. It is remarkable that MolAR is consistently faster than Gromacs, MDTraj and Pteros in the third test despite the fact that all of them share the same *xdrfile* C library for reading and writing xtc trajectories.

It is apparent that MolAR is very fast despite the fact that it is not optimized for performance except the trivial elimination of the redundant bound checks and the usage of the Rust optimizing compiler. In our tests MolAR is even faster than its logical predecessor Pteros, which was the fastest trajectory analysis library for the last decade ^12^. It is also clearly seen that the most popular trajectory analysis libraries written in Python, such as MDAnalysis and MDTraj, are also the slowest, lagging behind the natively compiled libraries even for the very basic IO-dominated tasks.

## Code Availability

MolAR is available for free under the Artistic License 2.0. Stable versions are published in the Rust community’s crate registry at https://crates.io/crates/molar. Latest development versions are available on GitHub https://github.com/yesint/molar. The code of all benchmarks described in this paper are available in the MolAR source code tree.

## Conclusions

In this work we presented MolAR -an experimental molecular analysis library for Rust programming language. We designed and implemented memory safe, fast and ergonomic atom selection abstractions in Rust, which are serving as universal building blocks for molecular analysis software and allow implementing efficient parallel analysis algorithms. In our benchmarks MolAR outperforms other popular molecular analysis libraries and tools, which makes it promising for developing computationally intensive analysis tasks.

## Acknowledgments

SY has received funding through the MSCA4Ukraine project 101101923, which is funded by the European Union. Views and opinions expressed are however those of the author(s) only and do not necessarily reflect those of the European Union. Neither the European Union nor the MSCA4Ukraine Consortium as a whole nor any individual member institutions of the MSCA4Ukraine Consortium can be held responsible for them.

## Conflicts of Interest

SY is an employee of Receptor.AI INC and has shares in Receptor.AI INC.

## References

1. Azevedo de Amorim, A., Hriţcu, C. & Pierce, B. C. The Meaning of Memory Safety. in Principles of Security and Trust (eds. Bauer, L. & Küsters, R.) 79–105 (Springer International Publishing, Cham, 2018). doi:10.1007/978-3-319-89722-6_4.

2. The Urgent Need for Memory Safety in Software Products | CISA. https://www.cisa.gov/news-events/news/urgent-need-memory-safety-software-products (2023).

3. Kevin Andrian Santoso, O., Kwee, C., Chua, W., Nabiilah, G. Z., & Rojali. Rust’s Memory Safety Model: An Evaluation of Its Effectiveness in Preventing Common Vulnerabilities. Procedia Comput. Sci. 227, 119–127 (2023).

4. Xu, H., Chen, Z., Sun, M., Zhou, Y. & Lyu, M. R. Memory-Safety Challenge Considered Solved? An In-Depth Study with All Rust CVEs. ACM Trans Softw Eng Methodol 31, 3:1-3:25 (2021).

5. Fourment, M. & Gillings, M. R. A comparison of common programming languages used in bioinformatics. BMC Bioinformatics 9, 82 (2008).

6. An Analysis of Scripting Languages for Research in Applied Computing. https://ieeexplore.ieee.org/abstract/document/6755355.

7. GlobalInterpreterLock - Python Wiki. https://wiki.python.org/moin/GlobalInterpreterLock.

8. Michaud-Agrawal, N., Denning, E. J., Woolf, T. B. & Beckstein, O. MDAnalysis: A toolkit for the analysis of molecular dynamics simulations. J. Comput. Chem. 32, 2319–2327 (2011).

9. MDAnalysis: A Python Package for the Rapid Analysis of Molecular Dynamics Simulations - SciPy Proceedings. https://proceedings.scipy.org/articles/Majora-629e541a-00e (2016).

10. McGibbon, R. T. et al. MDTraj: A Modern Open Library for the Analysis of Molecular Dynamics Trajectories. Biophys. J. 109, 1528–1532 (2015).

11. Yesylevskyy, S. O. Pteros: Fast and easy to use open-source C++ library for molecular analysis. J. Comput. Chem. 33, 1632–1636 (2012).

12. Yesylevskyy, S. O. Pteros 2.0: Evolution of the Fast Parallel Molecular Analysis Library for C++ and Python. (2015).

13. triomphe - crates.io: Rust Package Registry. https://crates.io/crates/triomphe (2024).

14. bindgen - crates.io: Rust Package Registry. https://crates.io/crates/bindgen (2024).

15. cmake - crates.io: Rust Package Registry. https://crates.io/crates/cmake (2024).

16. molar_molfile - crates.io: Rust Package Registry. https://crates.io/crates/molar_molfile (2024).

17. molar_xdrfile - crates.io: Rust Package Registry. https://crates.io/crates/molar_xdrfile (2024).

18. molar_gromacs - crates.io: Rust Package Registry. https://crates.io/crates/molar_gromacs (2024).

19. Klenin, K. V., Tristram, F., Strunk, T. & Wenzel, W. Derivatives of molecular surface area and volume: Simple and exact analytical formulas. J. Comput. Chem. 32, 2647–2653 (2011).

20. Abraham, M. J. et al. GROMACS: High performance molecular simulations through multi-level parallelism from laptops to supercomputers. SoftwareX 1–2, 19–25 (2015).

21. Humphrey, W., Dalke, A. & Schulten, K. VMD: Visual molecular dynamics. J. Mol. Graph. 14, 33–38 (1996).

